# From the inside out: An epibiotic *Bdellovibrio* predator with an expanded genomic complement

**DOI:** 10.1101/506055

**Authors:** Christoph M. Deeg, Tan T. Le, Matthias M. Zimmer, Curtis A. Suttle

## Abstract

Bdellovibrio and like organisms are abundant environmental predators of prokaryotes that show a diversity of predation strategies, ranging from intra-periplasmic to epibiotic predation. The novel epibiotic predator *Bdellovibrio qaytius* was isolated from a eutrophic freshwater pond in British Columbia, where it was a continual part of the microbial community. *Bdellovibrio qaytius* was found to preferentially prey on the beta-proteobacterium *Paraburkholderia fungorum.* Despite its epibiotic replication strategy, *B. qaytius* encodes a complex genomic complement more similar to periplasmic predators as well as several biosynthesis pathways not previously found in epibiotic predators. *Bdellovibrio qaytius* is representative of a widely distributed basal cluster within the genus *Bdellovibrio*, suggesting that epibiotic predation might be a common predation type in nature and ancestral to the genus.

## Introduction

Biotic factors regulating bacterial populations in nature are often assumed to be viral lysis and zooplankton grazing (1); however, an underappreciated cause of mortality is predation by other prokaryotes. Such predators, collectively referred to as Bdellovibrio and like organisms (BALOs), have evolved several times independently and deploy a variety of “hunting strategies”. Many facultative predators with broad host ranges, such as *Ensifer adhaerens* and *Myxococcus xanthus*, deploy a “wolfpack strategy” where a prey cell is surrounded by several predators and lysed (2, 3). Other, more specialized obligate predators have a narrower host range and specific predation strategies; for example, *Bdellovibrio spp.* enters the periplasm of the prey cell to consume the prey’s cytoplasm (4, 5).

*Bdellovibrio spp.* are delta-proteobacteria predators that use a biphasic lifestyle comprising an attack phase, in which a small, highly motile flagellated cell seeks out prey, and a growth phase, characterized by the predator penetrating the outer membrane of the prey cell and consuming its cytoplasm (5). During the growth phase, the predator forms a characteristic structure in the prey’s periplasm known as the bdelloplast, which consists of a rounded, osmotically stabile outer membrane of the prey cell and several replicating *Bdellovibrio* cells. The bdelloplast continues to grow until the resources of the prey cell are exhausted and culminates in the septation and release of several to dozens of new attack-phase cells. This dichotic lifestyle switch is mediated by a highly expressed riboswitch in *B. bacterivorous* (6). The related genera *Bacteriovorax* and *Predibacter* are in the family *Bacteriovoracea*, which is a sister family to the prototypical *Bdellovibrionacea* within the order *Bdellovibrionales* (7).

Curiously, the alpha-proteobacteria genus *Micavibrio,* which is unrelated to the *Bdellovibrionales* leads a remarkably similar lifestyle to *Bdellovibrio* species, with high prey specificity. However, these bacteria prey in an epibiotic fashion on the outside of the prey cell instead of penetrating into the periplasm (8). Due to their similar lifestyles, *Micavibrio* spp. are included into the BALOs.

Recently, an isolate of a newly described species, *Bdellovibrio exovorus*, in the family *Bdellovibrionacea* that is closely related to periplasmic bdelloplast-forming *Bdellovibrio* species, was shown to have an extremely narrow host range, and employ a different epibiotic replication strategy (9). In the attack-phase, cells of *Bdellovibrio exovorus* resemble those of other *Bdellovibrio* isolates; whereas, in growth-phase the cells do not penetrate into the cytoplasm, but stay attached to the outside of the prey, strongly resembling *Micavibrio* species. Further, in growth-phase, *B. exovorus* does not induce a bdelloplast and seems to extract the cytoplasmic contents of the prey across both membranes. Once the resources of the prey are exhausted, growth-phase result in binary fission releasing two progeny attack-phase cells. The comparatively small genome of *B. exovorus* has been linked to its epibiotic predation strategy and reductionist evolution from an ancestor capable of intra-periplasmic replication (9-11). *Bdellovibrio qaytius* is the second epibiotic predator within the genus *Bdellovibrio*. Its genomic complement, phylogenetic placement and environmental distribution broaden our understanding of the ecology and evolution of this genus.

## Materials and Methods

### Isolation and Culturing

An isolate of a lytic bacterium, here named *Bdellovibrio qaytius sp. nov.*, was obtained from a water sample collected near the sediment surface of a eutrophic pond in Nitobe Memorial Garden at the University of British Columbia, Canada (49°15′58″N, 123°15′34″W).As part of a bioassay for pathogens infecting heterotrophic protists a subsample of the water was inoculated into modified DY-V artificial freshwater medium with yeast extract and a wheat grain (12).

### Genome sequencing

For PacBio sequencing, exponentially growing mixed cultures containing *B. qaytius* as well as *Bodo saltans* NG1 and its virus BsV were centrifuged in a Sorvall SLC-6000 for 20 min and 5000 rpm at 4°C to remove eukaryotic cells (13). Particles in the supernatant concentrated approximately 100-fold by tangential flow ultrafiltration with at 30kDa cut-off (Vivaflow 200, PES) cartridge. To concentrate the cells further they were centrifuged at 28,000 rpm, 15°C for 8 h in a Beckman ultracentrifuge using a Ti90 fixed-angle rotor (Beckman-Coulter, Brea, California, USA), and then sedimented onto a 40% Optiprep 50 mM Tris-Cl, pH 8.0, 2mM MgCl_2_ cushion for 30 min at 28,000 rpm, and 15°C in a SW40Ti swing-out rotor. The Optiprep gradient was created by underlaying a 10% Optiprep solution in 50 mM Tris-Cl, pH 8.0, 2 mM MgCl_2_ with a 30% solution followed by a 50% solution and equilibration overnight at 4°C. One ml of concentrate from the 40% cushion was added atop the gradient and the concentrate was fractionated by centrifugation in an SW40 rotor for 4 h at 25000 rpm and 18°C. The fraction corresponding to the pathogen was extracted from the gradient with a syringe and washed twice with 50 mM Tris-Cl, pH 8.0, 2 mM MgCl_2_ followed by centrifugation in an SW40 rotor for 20 min at 7200 rpm and 18°C and were finally collected by centrifugation in an SW40 rotor for 30 min at 7800 rpm and 18°C. Purity of the concentrate was verified by fluorescence vs SSC of SYBR-Green stained samples (Invitrogen Carlsbad, California, USA) on a FACScalibur flow cytometer (Becton-Dickinson, Franklin Lakes, New Jersey, USA). High molecular weight genomic DNA was extracted using phenol-chloroform-chloroform extraction. Length and purity were confirmed by gel electrophoresis and by using a Bioanalyzer 2100 with the HS DNA kit (Agilent Technology). PacBio RSII 20kb sequencing was performed by the sequencing center of the University of Delaware. Reads were assembled using PacBio HGAP3 software with 20 kb seed reads resulting in a single contig of 3,376,027 bp, 97.08 x coverage, 99.92% called bases and a consensus concordance of 99.9954 % (14).

### Propagation and host range studies

Plaque assays were performed by mixing 0.5 ml putative host cultures in logarithmic growth stage and 10μl of Bdellovibrio qaytius stock culture with 4.5 ml molten 0.5% DY-V agar and incubation for 48h. Propagation of *Bdellovibrio qaytius* in liquid culture was monitored by PCR with custom primers set specific to *B. qaytius* 16S rDNA (Forward-5’-AGTCGAACGGGTAGCAATAC-3’, Reverse-5’-CTGACTTAGAAGCCCACCTAC-3’) as well as a BALO-specific primer set by Davidov et al. (7). To obtain a pure isolate of prey cells present in the mixed microbial assemblage, culture samples were streaked onto a DY-V agar plate and incubated at room temperature. Distinct colonies were picked and propagated in liquid DY-V medium. Propagation of *Bdellovibrio qaytius* using these cultures as hosts was confirmed by PCR. The identity of the prey cell cultures was confirmed by universal 16S rDNA Sanger sequencing (515F-5’-GTGYCAGCMGCCGCGGTAA-3’, 926R-5’-CCGYCAATTYMTTTRAGTTT-3’) (15). To clean up the predator culture, *E. coli* (Thermo Fisher) cells were grown in LB medium and pelleted at 3,900 x g (4500 rpm) for 10 min, washed with 10 ml of HEPES/CaCl2 buffer (25 mM HEPES, 2 mM CaCl2), centrifuged in a fixed angle rotor centrifuge at 3,900 x g for 5 min, and re-suspended in 19 ml of HEPES/CaCl2 buffer. This cell suspension was then inoculated with 1 ml of 0.8-μm PVDF membrane filtered lysate of the *Bdellovibrio* containing culture and *B. qaytius* propagation was monitored by PCR. Bdellovibrio remained viable at 4°C storage for up to two years and glycerol stocks of the native community containing *Bdellovibrio* as well as an inoculated E.coli TOP10 culture was stored at −80°C for archival purposes.

### Environmental sampling

The presence of *B. qaytius* in the Nitobe-Garden pond was determined in 20-L water samples that were taken bimonthly during spring and summer 2017 filtered through GF-A filters (Millipore, Bedford, MA, USA; nominal pore size 1.1 μm) laid over a 0.8-μm pore-size PES membrane (Sterlitech, Kent, WA, USA). The remaining particulate material was concentrated into 250 ml using a 30-kDa MW cut-off tangential flow filtration cartridge (Millipore, Bedford, MA, USA). DNA from these concentrates was extracted using phenol-chlorophorm extraction and subjected to PCR using *Bdellovibrio qaytius* specific 16S rDNA primers to confirm its presence.

### Microscopy

#### Negative staining transmission electron microscopy

Cultures of *Escherichia coli* TOP10 were inoculated with *B. qaytius* at two hour time intervals and infected cultures, as well as an uninfected control were diluted tenfold and fixed in 4% glutaraldehyde. Next, the samples were applied to the carbon side of formvar carbon-coated 400-mesh copper grids (TedPella, CA, USA) and incubated at 4°C in the dark overnight under high humidity. The liquid was then removed and the grids stained with 1% uranyl acetate for 30 s.

#### Ultra-thin sectioning transmission electron microscopy

For higher resolution images, cells of *E. coli* infected with *Bdellovibrio qaytius* were harvested at at 4h intervals, as well as from uninfected control cultures. Cells from 10 ml of culture were pelleted at 5000 xg in a Beckmann tabletop centrifuge using a fixed angle rotor. The pellet was resuspended in 0.2 M Na-cacodylate buffer, 0.2 M sucrose, 5% EM-grade glutaraldehyde, pH 7.4 and incubated for 2 h on ice. After washing in 0.2 M Na-cacodylate buffer, cells were post-fixed with 1% Osmium tetroxide. Samples were dehydrated through water/ethanol gradients and ethanol was substituted by acetone. Samples were embedded in an equal part mixture of Spurr’s and Gembed embedding and the resin was polymerized at 60°C overnight. Fifty-nm thin sections were prepared using a Diatome ultra 45° knife (Diatome, Switzerland) on an ultra-microtome. The sections were collected on a 400x copper grid and stained for 10 min in 2% aqueous uranyl acetate and 5 min in Reynold’s lead citrate. Image data were recorded on a Hitachi H7600 transmission electron microscope at 80 kV. Image J (RRID:SCR_003070) was used to compile all TEM images. Adjustments to contrast and brightness levels were applied equally to all parts of the image.

#### Fluorescence In Situ Hybridization Epifluorescence Microscopy

To confirm epibiotic predation, cultures for fluorescence in-situ hybridization (FISH) were prepared as outlined below. Two 10-ml volumes of *E. coli* TOP10 were centrifuged at 3900 xg in a Beckman tabletop fixed angle centrifuge (4500 rpm) for 10 min, washed with 5 ml of HEPES/CaCl2 buffer, centrifuged at 3900 g for 5 min, and re-suspended in 9 ml of HEPES/CaCl2 buffer. One ml of *B. qaytius* containing culture was added to the resuspended *E. coli* while another served as a control. Both cultures were incubated at room temperature for 24 h and were centrifuged again at 3900 x g (4500 rpm) for 10 min, washed with 10 ml of PBS, centrifuged at 3,900 x g for 5 min, and re-suspended in 5 ml of PBS. Two ml of the cultures were fixed in a 1:3 dilution of 10% buffered formalin (pH 7.0; 10 ml of 37% formaldehyde, 0.65 g NalJHPOlJ, 0.4 g NaH_2_PO_4_. 90 ml of Milli-QTM H2O) at 4°C for 3 h. Cells were then centrifuged again at 3,900 x g, washed twice in 10 ml of PBS, re-suspended in 10 ml of a mixture of PBS and 96% EtOH (1:1), and vortexed. In order to localize the predator an Alexa-488 tagged probe specific to *Bdellovibrio* 16S rDNA was designed (5’-/5Alexa488N/TGCTGCCTCCCGTAGGAGT-3’) based on Mahmoud et al. which also served as a template for the incubation protocol (16). Ten μl of sample was spotted onto a 70% EtOH-cleaned slide, dried at room temperature, and then taken through a dehydration series of 50%, 80%, and 95% EtOH. 25 μl of the hybridization master mix (20 mM Tris-HCl [pH 7.4], 0.1 % SDS, 5 mM EDTA, 0.8 M NaCl, 37% formalin, 1 ng/μl of final probe concentration) was added onto the sample. A cover glass was placed onto each sample and the slides incubated for 2 h at 46°C. With the cover slip removed, the slides were subsequently submerged into a bath of wash buffer and incubated at 48°C for 30 min. Slides were rinsed with sterile deionized H2O and dried at room temperature. A drop of ProLongTM Diamond Antifade Mountant with DAPI (4,6-diamidine-2-phenylindole) was spotted onto a new cover glass and placed on the sample. Finally, the slides were incubated at room temperature in the dark for 24 h prior to observation on an Olympus FV 1000 system.

### Annotation

The genome was circular and 3,348,710 bp in length. Genome annotation was performed using the automated NCBI Prokaryotic Genome Annotation Pipeline (PGAAP). In parallel, open reading frames were predicted using GLIMMER (RRID:SCR_011931) with default settings (17). Translated proteins were analyzed using BLASTp, CDD RPS-BLAST and pfam HMMER. These results were used to refine the PGAAP annotation. Signal peptides and trans-membrane domains were predicted using Phobius (18). The annotated genome is available under the accession number CP025544. Metabolic pathways were predicted using the Kyoto Encyclopedia of Genes and Genomes (KEGG RRID:SCR_012773) automatic annotation server KAAS and Pathway Tools (RRID:SCR_013786) (19, 20).

### Phylogenetic analysis

Full length 16S rDNA sequences of completely sequenced isoaltes of *Bdellovibrio* spp., as well as full-length uncultured top BLAST hits were downloaded from NCBI. Alignments of rDNA sequences were performed in Geneious R9 (RRID:SCR_010519) using MUSCLE with default parameters (RRID:SCR_011812)(21). Maximum likelihood trees were constructed with RAxML ML search with 1000 rapid bootstraps using GTR+GAMMA (22).

Phylogenetic analysis of the genome content by orthologous gene clusters was performed by OrthoMCL (RRID:SCR_007839) (23) using whole genome sequences downloaded from NCBI. OrthoMCL was run with standard parameters (Blast E-value cutoff = 10-5 and mcl inflation factor = 1.5) on all protein-coding genes of length ≥ 100 aa. This resulted in the definition of 4242 distinct gene clusters.

## Results

### Isolation, host range and distribution

A lytic pathogen of bacteria was collected in a mixed microbial assemblage from a temperate eutrophic pond in southwestern British Columbia, Canada. Based on full-genome sequencing and electron microscopy of the infection cycle the pathogen was determined to be a new species of bacterium, here named *Candidatus Bdellovibrio qaytius (*subsequently referred to as *B. qaytius*), after “qLa:yt” (“kill it”) in hən □q□əmin□əm□ the language of the Musqueam tribe of Coast Salish, the indigenous peoples from who’s territory it was isolated. *B. qaytius* propagated in a mixed microbial assemblage from the sample site (Supplementary Figure 1). Additionally, *B. qaytius* could also be propagated on a specific isolates from this assemblage that was identified as the beta-proteobacterium *Paraburkholderia fungorum* where high numbers of putative attack-phase *B. qaytius* cells could be observed under phase-contrast microscopy (Supplementary Figure 1). Inoculation of *B. qaytius* in cultures of *Pseudomonas fluorescence*, another isolate from the native mixed assembly, or *E. coli* resulted in the observation of weak PCR signals after propagation, presumable due to carry-over(Supplementary Figure 1). Similarly, small clear plaques were only observed on plates of *P. fungorum,* but not on plates of *Pseudomonas fluorescence*, or *E. coli (*Figure 1, Supplementary Figure 2). *Bdellovibrio qaytius* was detected at several time points in DNA extracted from water concentrates from Nitobe Gardens UBC in 2017, four years after the initial isolation, indicating the population persists in the pond (Supplementary Figure 3).

**Figure 1:**
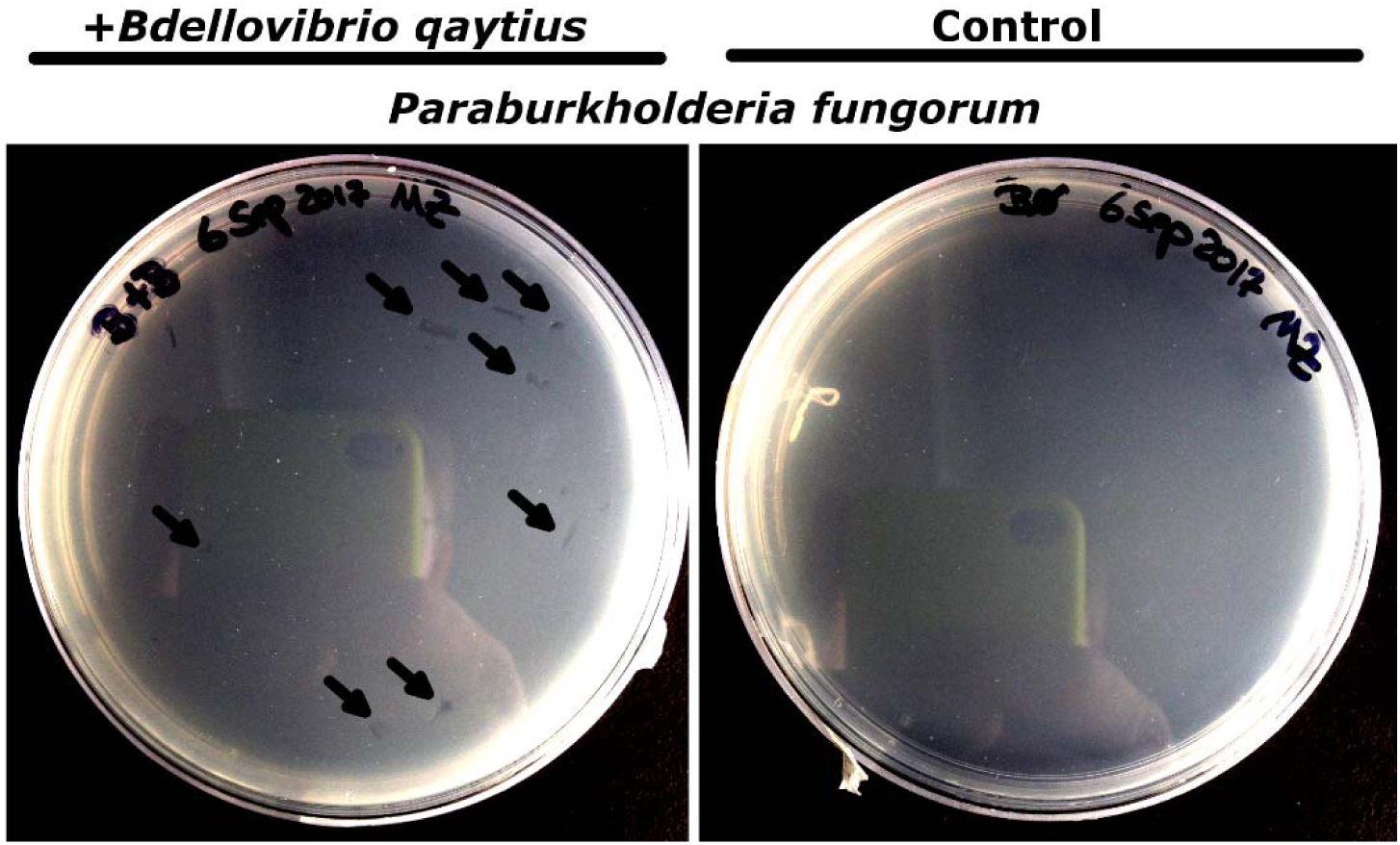
Plaque assay. *Bdellovibrio qaytius* added to plates of *Paraburkholderia fungorum* on the left, untreated control on the right. Arrow indicate location of plaques 24hpi.

### Morphology and replication cycle

*Bdellovibrio qaytius* attack-phase cells are free swimming highly motile flagellated rods that are about 1 μm by 0.4 μm in size and exhibit a sheathed flagellum (Figure 2 A). Once attack-phase cells contact a prey cell, they attach irreversibly and form a broad predatory synapse and discard the flagellum (Figure 2 B). No invasion of the prey cell was observed during growth-phase, nor were bdelloplasts, implying that *Bdellovibrio qaytius* is an epibiotic predator (Figure 2). The growth-phase cell attached to the prey cell empties the cytoplasm of the host cell, leaving behind an empty ghost cell (Figure 2 C,E,G). Simultaneously, the growth-phase *Bdellovibrio* cell grows in size and once the resources of the prey cell are exhausted, the growth-phase cumulates in binary fission and the production of two offspring attack-phase cells that repeat actively searching for new prey cells by rapid locomotion. Throughout the growth-phase, the cell membranes of the prey as well as the predator remain intact and instead of periplasmic invasion, an electron dense layer is observed on both the prey’s and predator’s membranes suggesting that a high concentration of effector molecules like transmembrane transporters are likely recruited to these sited to facilitate predation (Figure 2 F). Predation was dependent on the growth-phase of the host cell with cells in logarithmic growth supporting the highest *B. qaytius* concentrations.

**Figure 2:**
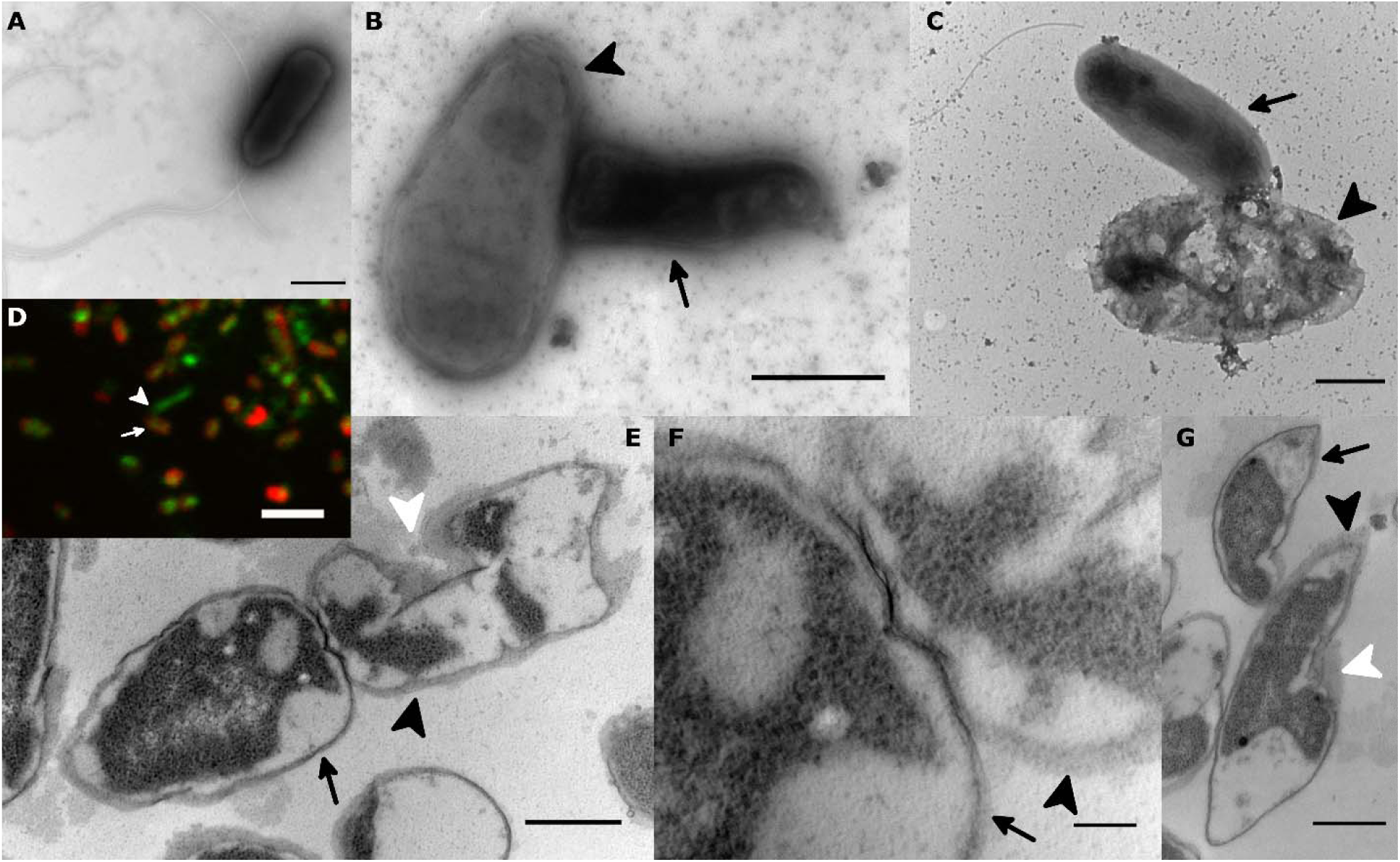
Bdellovibrio qaytius predation strategy and replication. *A*: Negative staining electron micrograph of an attack-phase cell showing the characteristic sheathed flagellum. B: Negative staining electron micrograph of an early growth-phase cell (arrow) attaching to a prey cell (arrow head) with a broad predatory synapse. C: Negative staining electron micrograph of a late growth-phase *Bdellovibrio qaytius* (arrow) next to a ghost cell of a prey cell (arrow head). The flagellum is detached in preparation for binary fission. D: FISH epifluorescence micrograph of *B. qaytiu* (red with specific FISH probe, arrow) attached to host cell (green DAPI stained, arrow head). E: Thin-section electron micrograph of growth-phase *B. qaytius* (black arrow) attached to a prey cell (black arrow head). The prey cell has an emptied cytoplasm and shows an invagination of the membrane (white arrow head).F: Thin section micrograph close-up of the predatory synapse shown in E. The membrane of the predator (arrow) and the prey cell (arrow head) remain intact, but show electron dense signatures. G: growth-phase *B. qaytius* (black arrow) showing polar attachment to a prey cell (black arrow head) that also shows an invaginating cell membrane (white arrow head). Scale bar in D: 2.5 μm, other scale bars + 500 nm.

### *Bdellovibrio qaytius* has a complex genome for an epibiotic predator

#### Genome structure and content

The 3,348,710-bp *B. qaytius* genome is similar in size to periplasmic *Bdellovibrio* spp., but considerably larger than that of *Bdellovibrio exovorus*, another epibiotic predator with a 2.66 Mb genome and with 38.9% also has the lowest GC content of any species within the genus. The GC content is relatively constant and exhibits a dichotomy in GC-skew that is typical of a circular bacterial genome (Figure 3A). Furthermore, the *B. qaytius* genome contains one complete rDNA operon, similar to other epibiotic predators, and 31 tRNAS, three non-coding RNAs (ssrS, rnpB and ffs), and three putative riboswitches (Figure 3B). A total of 3166 protein coding genes were identified and are distributed equally between the plus and minus strands(Figure 3A). These proteins represent 22 different functional clusters of orthologous genes (Figure 3B).

**Figure 3:**
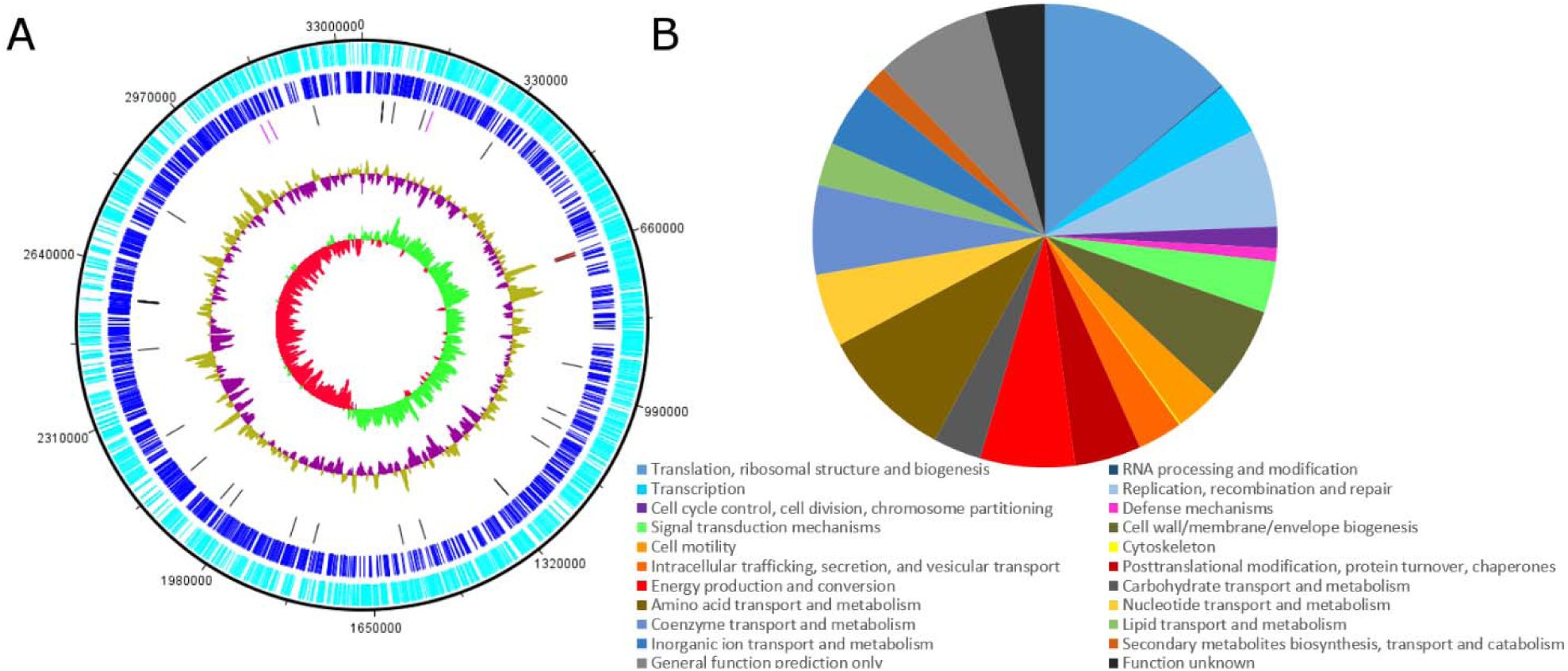
Bdellovibrio qaytius *genome.* A: Genomic map of *B. qaytius*. From outside to inwards: Plus-strand CDS (light blue), Minus-strand CDS (dark blue), tRNAs (black), rRNAs (red), and non-coding RNAs (pink), GC-content (purple/mustard), and GC-skew (red/green). B: Abundance of 866 identified functional clusters of orthologous genes in the *B. qaytius* genome

#### Metabolism

The *B. qaytius* genome encodes a metabolism typical for a predatory bacterium. Glycolysis and the complete TCA cycle, as well as a core set of pentose phosphate pathway genes are present, suggesting *B. qaytius* is capable of several sugar conversions, and is able to provide the precursors for riboflavin biosynthesis. Pyruvate metabolism is coded for, but propanoate metabolism is only partially possible. A vitamin B6 biosynthesis pathway is encoded and acetyl-CoA biosynthesis is possible via pantoate. Nicotinamide metabolism and biosynthesi pathways also exist. Oxidative phosphorylation is encoded with the exception of cytochrome C reductase. The presence of core mevalonate pathway enzymes suggests this pathway i functional.

Based on inferred CDS, *B. qaytius* can synthesize pyrimidines and purines *de novo*, and in contrast to other epibiotic predators, produce inosine. A complete DNA polymerase complex facilitates DNA replication. As well, all types of DNA repair pathways are present, including base excision, nucleotide excision, mismatch repair and homologous recombination, the latter being limited to single-stand-break repair.

*Bdellovibrio qaytius* encodes a complete ribosome except for the non-essential protein L28, and there is a complete set of tRNAs loaded by aminoacyl-tRNA-synthetases for every amino acid. The genome encodes a core RNA degradasome for post-transcriptional regulation and nucleotide recycling. Amino-acid biosynthesis pathways are limited and only cysteine, methionine, glutamate, lysine, proline and threonine can be completely synthesized *de novo*. Glycine and serine can be synthesized via the one-carbon pool pathway using tetrahydrofolate. Aspartate, alanine leucine, isoleucine, valine phenylalanine and tyrosine can be converted from their direct precursors, which are presumably acquired from the prey.

Complete fatty-acid degradation pathways are coded for, but fatty-acid elongation seems limited. Also encoded are complete sec and gsp pathways for protein secretion, as well as a partial tat pathway (subunits tatA, tatC, tatD). Extensive peptidoglycan production, as well as partial lipopolysaccharide biosynthesis pathways putatively decorate the periplasmic space and cell surface.

### Regulatory elements

Master regulators, such as sigma factor 28 / FliA, which is proposed to enable the switch between attack and growth-phase modes in other *Bdellovibrio* species, is present in *B. qaytius* and might work alongside putative riboswitch elements similar to those found in *B. bacterivorus* (6). No homologue of the host interaction (“hit”) locus protein bd0108, of *B. bacterivorus*, was found in *B. qaytius*, suggesting that it may deploy a different pilus regulation mechanism, and therefore might not be able to switch between facultative and obligate predation.

### Predatory arsenal

The *B. qaytius* genome shows many adaptations to a predatory lifestyle. A complete biosynthesis pathway for flagellar assembly and regulation provides locomotion in the attack-phase. Attack phase cells are likely guided by a canonical, almost, complete chemotaxis pathway that is only missing cheY. Substrate recognition is putatively mediated by tatC and two copies of von Willebrand factors (24). A type IV pilus appears to be present that is putatively involved in prey-cell attachment through pilZ. To access resources within the prey, an array of transporters are coded for, many of which show signal peptides that facilitate export, and might insert into the prey-cell membrane. ABC-type transporters likely import phosphate (pst), phosphonate (phn), and lipopolysaccharides (lpt), while there appears to be partial ABC transporter systems for lipoproteins, thiamine, branched-chain amino acids, oligo and dipeptides, microcin, phospholipids, biotin, daunorubiscine, alkylophaosphate, methionine, iron and siderophores, cobalt, sugar and organic solvents. Non-ABC transporter system CDS are present for potassium (kdp), biopolymers (exbD and tol), iron (ofeT), heavy metals (cusA), biotin (bioY), threonine (rhtB), as well as for several multidrug exporters (bcr, cflA, arcB). CDS for low-specificity transporters include MFS and EamE transporters, as well as transporter for ions and cations, macrolide, chromate, as well as sodium-dependent transporters and others of uncharacterized specificity.

### *Bdellovibrio qaytius* is a basal representative of its genus

In phylogenetic analysis of the 16S rDNA locus, *B. qaytius* occupies a well-supported basal branch within the genus *Bdellovibrio*; the closest relatives are found in environmental amplicon sequences, as well as in poorly characterized isolates (Figure 4A). Notably, these strains from a tight cluster basal to *B. exovorus* and *B. bacterivorus* strains and show rather low boot-strap support within their clade despite being closely related. This cluster appears to be equivalent to the “cluster 2” described by Davidov et al. (7).

**Figure 4:**
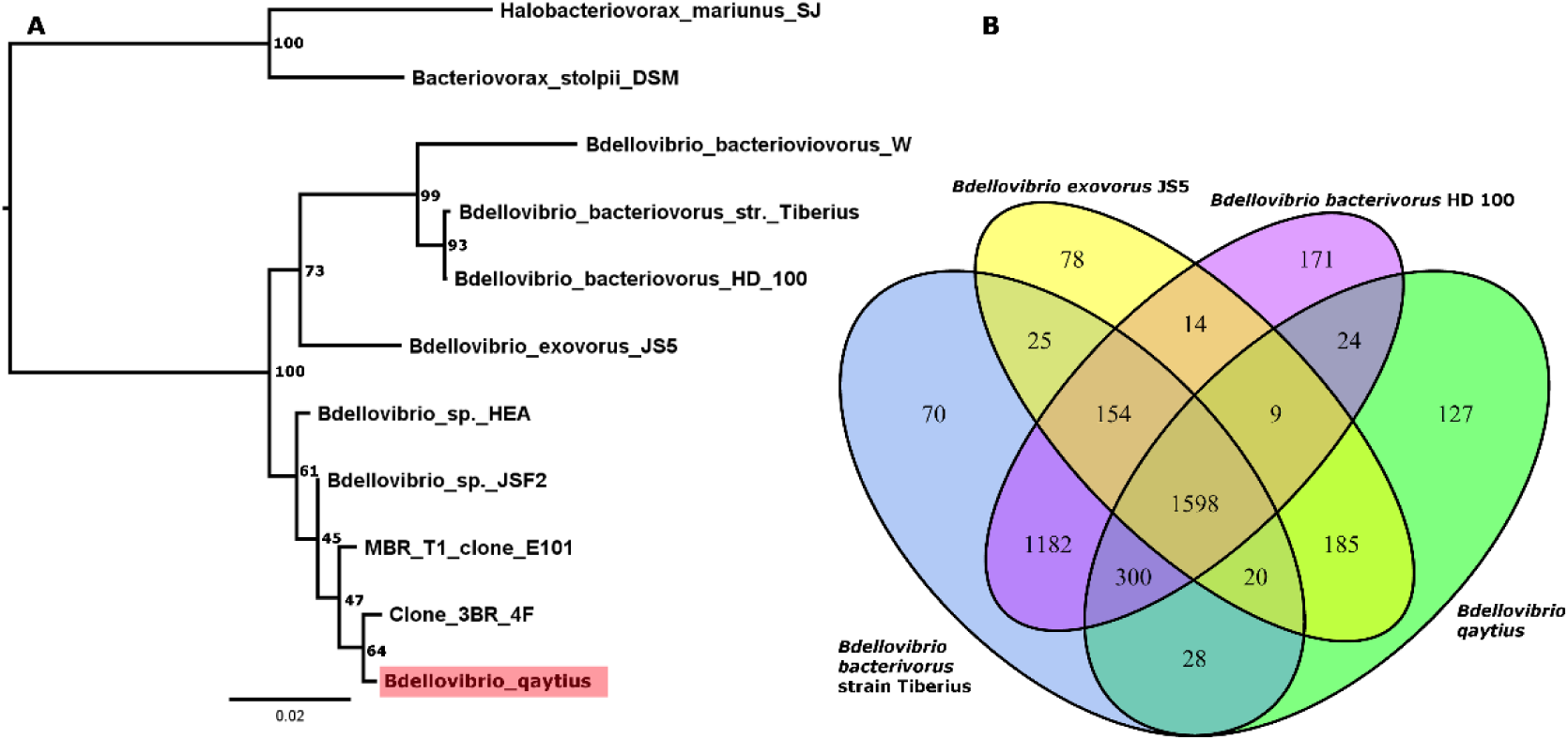
Bdellovibrio qaytius *phylogenetic placement.* A: 16S phylogenetic tree showing th completely sequenced *Bdellovibrio* species as well as the top BLAST hits to *Bdellovibrio qaytiu* from uncharacterized isolates, as well as from metagenomic data. B: Shared gene cluster analysi of complete *Bdellovibrio* genomes comparing epibiotic and periplasmic species.

Shared gene cluster analysis was congruent with the 16S phylogeny with the majority of gene clusters in *B. qaytius* being shared with other members of the genus *Bdellovibrio* (Figure 4B). Surprisingly, *B. qaytius* shares more than 300 gene clusters with periplasmic predators, which are not found in the epibiotic predator *B. exovorus*, despite their similar predation strategy. On the other hand, the epibiotic predators *B. qaytius* and *B. exovorus* shared more than 130 genes that are not found in periplasmic predators, and might be involved in epibiotic predation. These exclusively epibiotic genes within members of the genus *Bdellovibrio* include CDS for proteases, peroxiredoxin, glutathione-dependent formaldehyde-activating enzyme, nucleases, hydrolases thioesterase, polysaccharide deacetylase, an amino-acid ABC transporter, as well as many poorly characterized proteins. Gene clusters exclusively shared with *B. exovorus*, as well as the phylogenetically distant *M. aeruginosavorus* that also deploys an epibiotic predation strategy, are remarkable as they might highlight a common set of genes are required for epibiotic predation. Six such genes that were identified include anhydro-N-acetylmuramic acid kinase, peptidoglycan translocase, a FAD/NAD binding protein, a cation-transporting P-type ATPase and a pseudouridine synthase.

## Discussion

### *Bdellovibrio qaytius* is an epibiotic predator of the beta-proteobacterium *Paraburkholderia fungorum*

While *B. qaytius* 16S rDNA was detected by PCR in liquid cultures of *Pseudomonas fluorescence* as well as *E. coli* after two rounds of propagation, a weak signal compared to cultures containing *Paraburkholderia fungorum* as prey suggests the passive carry-over of *B. qaytius* cells or genetic material in the absence of replication (Supplementary Figure 1). This is in line with the higher density of putative attack-phase *B. quaytius* cells observed in *P. fungorum* cultures compared to the other putative hosts. Ultimately, plaque assay confirmed the beta-proteobacterium *Paraburkholderia fungorum* as the only host organism of *B. qaytius* observed in the present study.

### Despite an epibiotic phenotype, *Bdellovibrio qaytius* shares many genes with periplasmic predators

Microscopic analysis clearly shows *B. qaytius* deploying an epibiotic predation strategy, which is reflected in its genome that shares several features previously identified to be involved in epibiotic predators such as *B. exovorus* and *Micavibrio aeruginosavorus*. These include physical features such as the number of rDNA loci, as well as metabolic capabilities based on gene content that suggests limited fatty-acid elongation and the absence of polyhydroxyalkanoate depolymerase and the siderophore aerobactin, all present in periplasmic predators (11). In contrast, *B. qaytius* also has coding sequences for the biosynthesis of isoleucine and tyrosine, as well as for riboflavin and vitamin B6, which had been found in periplasmic but not epibiotic predators reflecting its comparatively large and complex genome (10). The linkage of these genes with periplasmic replication was by association and not for functional reasons; hence, the presence of several of these genes in *B. qaytius* may simply reflect its relatively larger genome size. Cluster analysis of orthologous genes in *B. qaytius,* other *Bdellovibrio spp.* as well as the unrelated BALOs *Micavibrio aeruginosavorus* and Halobacteriovorax marinus reveals just six gene clusters associated with epibiotic predation. Strikingly, these genes suggest that the prey peptidoglycan is salvaged by N-acetylmuramic-acid kinase as well as a peptidoglycan translocase that is specific to epibiotic predators. Since this limited complement of genes was found between distantly related taxa, there might not be a clear functional separation between epibiotic and periplasmic predators. This is consistent with epibiotic predation evolving independently within the genera *Bdellovibrio* and *Micavibrio*. Therefore, conclusions regarding function based on gene-cluster analysis should be interpreted with caution, especially since functionally equivalent proteins can group into different gene clusters, and thus may escape such analysis. Accordingly, gene-cluster comparison among species in the same genus that deploy different predation strategies is more informative and revealed a specialized complement of proteases nucleases, hydrolases and detoxifying enzymes in epibiotic *Bdellovibrio* species (Figure 4 B). Notably, the addition of *B. qaytius* as a second epibiotic *Bdellovibrio* species greatly decreased the number of genes associated with epibiotic predation, as its large genome has greater overlap with periplasmic *Bdellovibio* species. Further, this suggests that different mechanisms can be deployed in both, epibiotic and periplasmic predation.

### Epibiotic predation is a common strategy of environmental *Bdellovibrio* species and could be the ancestral phenotype

The closest known relatives to *B. qaytius* are poorly characterized isolates, and environmental 16S rRNA sequences from diverse environments. These environments include “commercial aquaculture preparations”, soils, waste-water activated sludge, and iron-oxidizing freshwater environments (7, 25-27). This broad diversity of habitats suggests that *Bdellovibrio* species that are closely related to *B. qaytius* are widely distributed in freshwaters, and could be a major contributor to global BALO diversity. Because of the phylogenetic placement and the broad distribution of related isolates, epibiotic predation might be common among BALOs, despite being underrepresented in isolates. The recurrent detection of populations of *B. qaytius* in the pond from which it was isolated confirms that is part if the natural community and therefore supports the idea that epibiotic predation might be common in nature. Additionally, both *B. qaytius* and *B. exovorus* are epibiotic and branch basal within the genus, compared to the periplasmic predators. This suggests that epibiotic predation might be the ancestral predation type in the genus.

Detailed knowledge of environmental BALO diversity is still missing. The discovery and analysis of *Bdellovibrio qaytius* provides new insights into this fascinating group. Its intermediate genomic complement blurs the lines between what is required for epibiotic and periplasmic replication. Moreover, as a representative of a basal and widespread member within the genus, *Bdellovibrio*, it suggests that epibiotic predation is common in the environment, and might be the ancestral form of predation in this genus.

## Acknowledgements

The work was supported by grants to C. S. from the Natural Sciences and Engineering Research Council of Canada (NSERC; 05896), Canada Foundation for Innovation (25412), British Columbia Knowledge Development Fund, and the Canadian Institute for Advanced Research (IMB). C. D. was supported in part by a fellowship from the German Academic Exchange Service (DAAD). T. L. was supported in part by an award by the Natural Sciences and Engineering Research Council of Canada. The authors would like to thank Jill Campbell, the coordinator for the Musqueam Language and Culture Department (https://fnel.arts.ubc.ca/community/musqueam-nation/musqueam-language-and-culture/), for her guidance in identifying a suitable species name based on the hən □q□əmin□əm□ language.

## Supplementary information

**Supplementary Figure 1:**
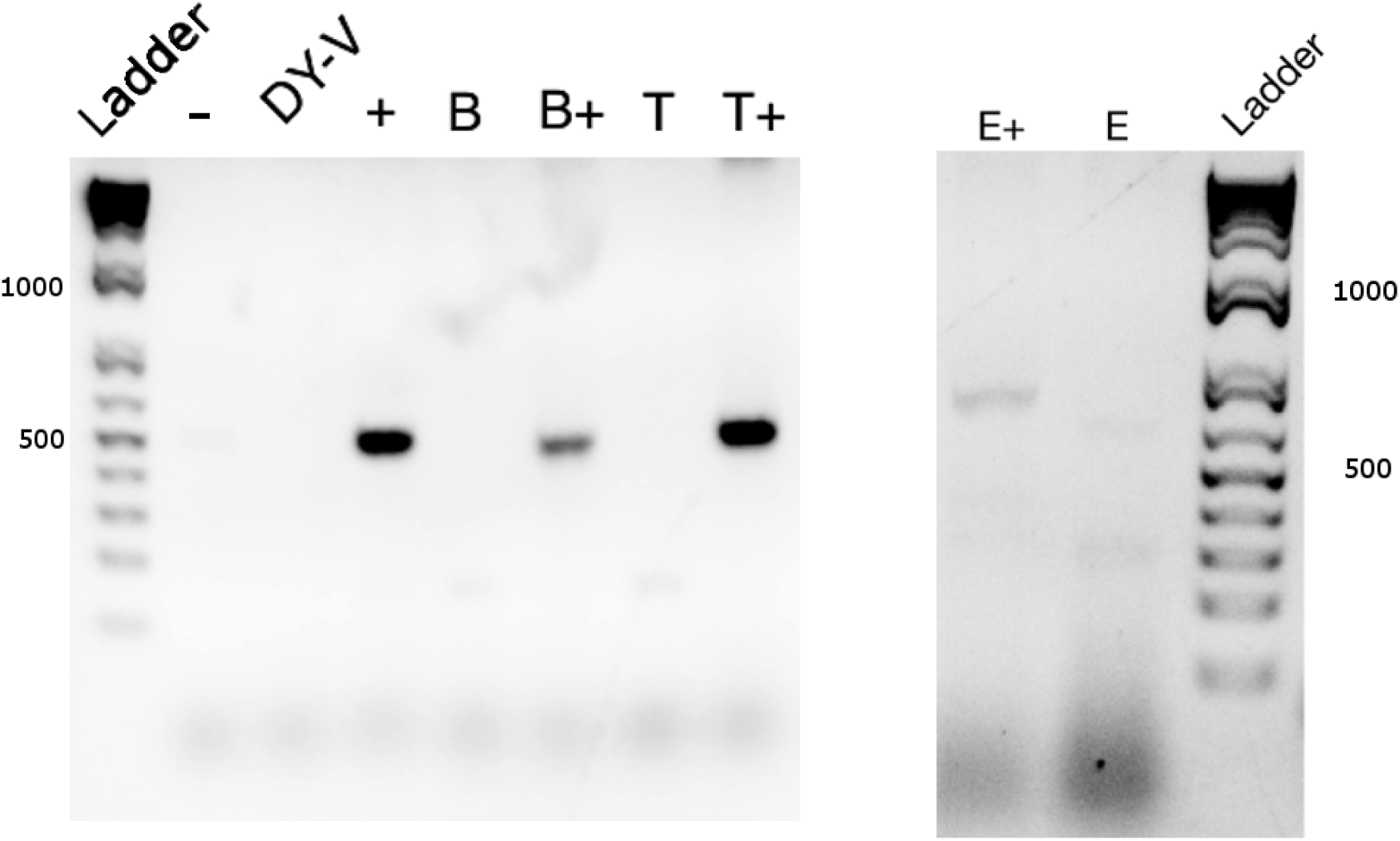
Bdellovibrio qaytius host range.B. qaytius propagation was assessed by PCR using strain-specific primers: -= water control, DY-V=medium control, += Positive control from mixed Nitobe garden pond assemblage culture, B= *Pseudomonas fluorescence* culture, B+= *Pseudomonas fluorescence* culture after inoculation and two propagation cycles showing carry over effects of *B. qaytius*, T= *Paraburkholderia fungorum* culture, T+ = *Paraburkholderia fungorum* culture after inoculation and two times propagation of *B. qaytius* showing active replication. E+ = *E. coli* culture after inoculation and two propagation cycles showing carry over effects of *B. qaytius* E = *E. coli* culture (E+ and E assessed with BALO primers [7]).

**Supplementary Figure 2:**
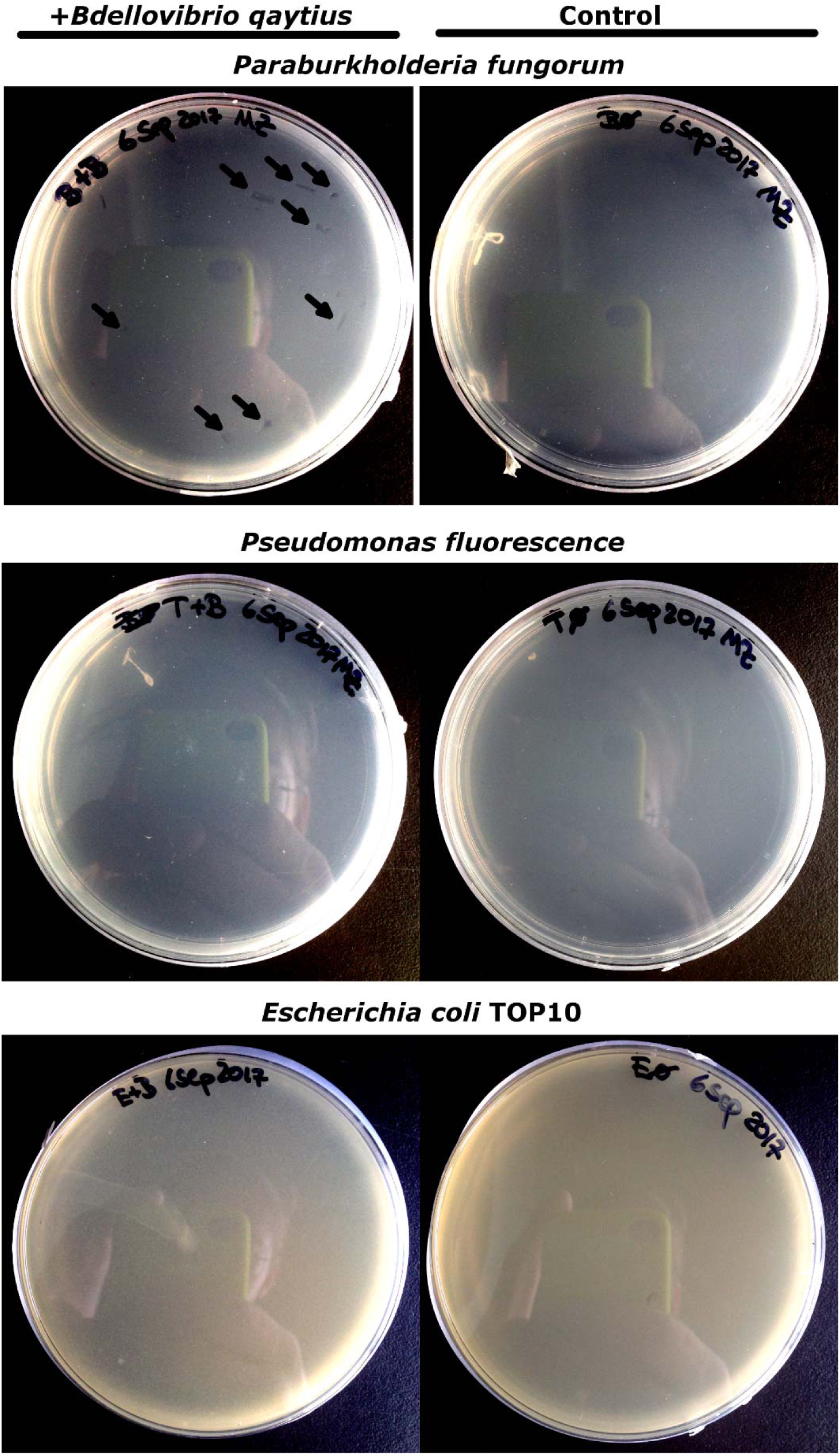
Bdellovibrio qaytius plaque assay. Left column shows *B. qaytiu* infected plates, right column shows the control. Plaques highlighted by arrows.

**Supplementary Figure 3:**
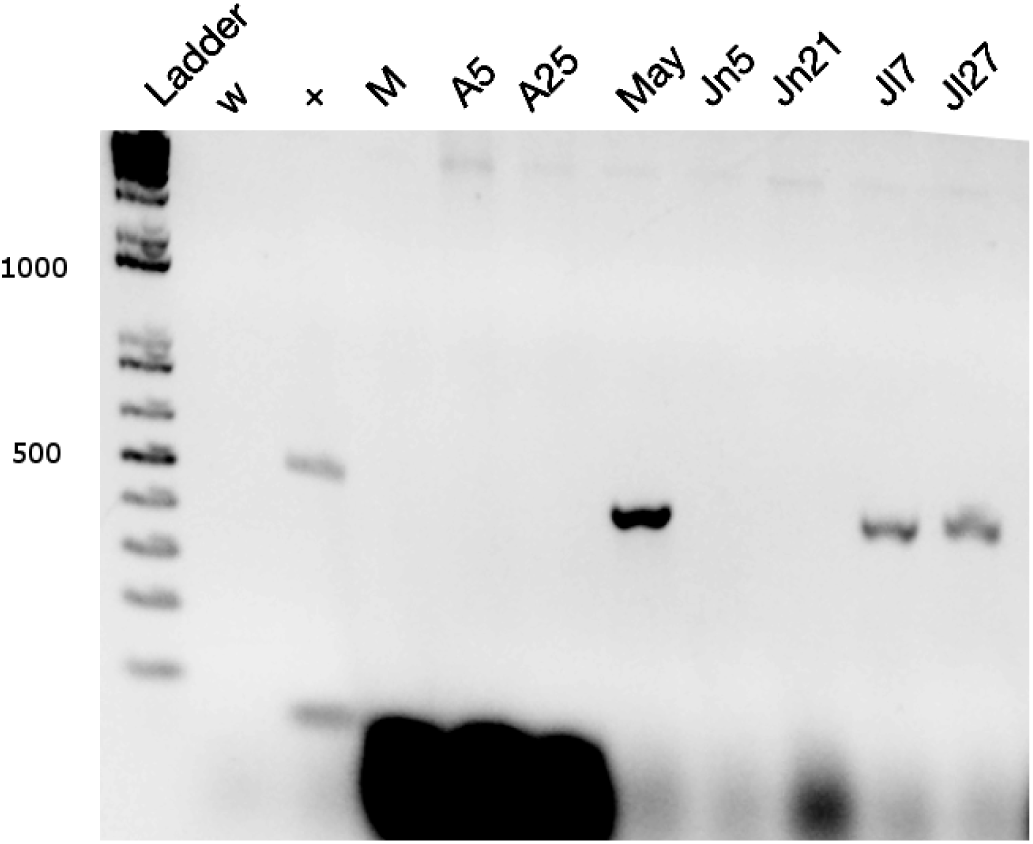
Bdellovibrio qaytius continued presence in Nitobe Gardens pond.: *Bdellovibrio qaytius* detection at various time points with *B. qaytius* specific primers in sample collected from Nitobe gardens UBC: w= water control, += positive control, M= 23 March 2017, A5= 5 April 2017, A25= 25 April 2017, May= 15 May 2017,Jn5= 5 June 2017, Jn21= 21 June 2017, Jl7= 7 July 2017, Jl27= 27 July 2017.

